# A missense mutation in *FBXL3* links circadian regulation to out-of-season estrus in sheep

**DOI:** 10.64898/2026.06.18.733072

**Authors:** Yang Yang, Na Zhang, Tingting Li, Haitao Wang, Xun Huang, Runlin Ma, Haoran Zhang, Xinxiao Jing, Ran Di, Qing Xia, Xiaoyun He, Xiaofei Guo, Xiaosheng Zhang, Yu Jiang, Ran Li, Mingxing Chu, Qiuyue Liu

## Abstract

Seasonal breeding is a remarkable adaptive trait, but it constrains efficient production in the sheep industry. Recent studies have shown that seasonal breeding is associated with endogenous circannual rhythms, which are regulated in part by the circadian clock system. FBXL3, a pivotal component of the SCF (SKP1 – CUL1 – F-box) E3 ubiquitin ligase complex, is a known determinant of the mammalian circadian period. In this study, we identified a missense mutation, T183M, in FBXL3 through selective sweep analysis. The allele frequency of this mutation differed significantly between sheep breeds exhibiting year-round estrus and those showing seasonal breeding patterns. The association between the T183M mutation and seasonal breeding was further validated using an ovariectomized, estradiol-implanted sheep model. We then generated mice carrying the homologous T183M mutation and found that they exhibited significantly lengthened circadian periods, accompanied by reduced CRY1 expression and increased CLOCK expression. Co-immunoprecipitation assays confirmed that the mutation reduced the interaction between FBXL3 and CRY1. These findings demonstrate an evolutionarily conserved role of FBXL3 in the circadian clock system. We propose that the T183M mutation disrupts day-length recognition, thereby influencing seasonal estrus in sheep.

**Author summary:** Seasonal breeding limits sheep productivity and is regulated by circadian rhythms, yet the key genetic determinants remain poorly understood. Here, we identified an *FBXL3* T183M missense mutation whose allele frequency differed markedly between year-round-estrous and seasonally breeding sheep, and validated its association with seasonal reproduction in a sheep population. Functional analyses in mutant mice showed that this variant lengthened the circadian period, disrupted the expression of core clock genes, and weakened the interaction between FBXL3 and CRY1. These findings suggest that the *FBXL3* T183M variant impairs day-length perception, thereby modulating seasonal estrus in sheep. Our study reveals a conserved circadian mechanism underlying seasonal breeding and highlights *FBXL3* T183M as a promising genetic target for improving reproductive performance in sheep.

## 1 Introduction

The Earth is subject to periodic environmental changes, such as the diurnal cycle driven by the Earth’s rotation and the seasonal cycle resulting from its revolution around the Sun. To cope with these rhythmic environmental fluctuations, organisms have evolved adaptive physiological and behavioral strategies, including reproduction, hibernation, and molting. Among these regulatory cues, the photoperiod, defined as the alternation of light and darkness within a 24-hour cycle and typically represented by the length of daylight, is regarded as the most reliable seasonal signal due to its consistent annual periodicity. Because other environmental factors such as temperature and food availability can fluctuate unpredictably, photoperiod provides a stable and anticipatory cue that allows organisms to align their biological processes with seasonal changes [1]. In regions outside the tropics, photoperiod varies predictably with the seasons. To ensure that offspring are produced during periods most favorable for survival, many animals have evolved mechanisms that use photoperiod as an endogenous “calendar” to regulate reproductive timing, a phenomenon known as photoperiodism [2]. Accordingly, animals can be classified according to their reproductive responses to day length as long-day breeders, such as hamsters and many avian species, or short-day breeders, such as most goats, sheep, and deer.

It is now well-established that the principal physiological pathway by which light regulates seasonal estrus in mammals involves the hypothalamic-pituitary-gonadal (HPG) axis. Light signals are perceived by the retina and converted into optic nerve impulses, which are transmitted to the suprachiasmatic nucleus (SCN) of the hypothalamus [3, 4]. As the master biological clock in mammals, the SCN relays photoperiodic information to the pineal gland, thereby altering the duration of nocturnal melatonin (MEL) secretion [5, 6]. The pars tuberalis (PT), which expresses melatonin receptor 1 (MT1), acts as a major target tissue for MEL in response to changes in melatonin signaling [7, 8]. Under the regulation of transcription factors and coactivators such as TEF, EYA3, and SIX1[9], long-day photoperiods increase the secretion of thyrotropin (TSH), which subsequently acts on adjacent hypothalamic ependymal cells, particularly tanycytes [10]. This process upregulates type II deiodinase (DIO2) expression in the mediobasal hypothalamus (MBH), while downregulating type III deiodinase (DIO3) expression [11–13]. Short-day photoperiods exert the opposite effect. These changes ultimately modulate the local conversion of thyroxine (T4) to triiodothyronine (T3) in the MBH, thereby influencing the pulsatile release of gonadotropin-releasing hormone (GnRH) from the hypothalamus. Through positive and negative feedback loops of reproductive hormones within the HPG axis, this pathway regulates the transition between estrus and anestrus [12, 14]. Currently well-defined targets involved in the regulation of seasonal estrus include the following: 1) the PT, which functions as a circannual molecular clock for estrous traits and responds to changes in MEL, while the expression of related genes in this tissue is not affected by downstream sex hormone fluctuations; 2) tanycytes, which serve as local T3-generating cells and in which estrogen and progesterone regulate the expression of specific genes to initiate the transition from estrus to anestrus; and 3) hypothalamic Kiss1 and Npvf neurons, which act as downstream output regulators and are modulated by ovarian sex hormones. Among these tissues and cell types, the PT has been identified as a key upstream regulator and circannual timer governing seasonal estrus [11, 15, 16]. However, the precise mechanism by which photoperiodic signals are transmitted from the circadian clock system to the HPG axis, and whether specific intermediate proteins mediate this regulatory process, remains unclear. In 2007, several research groups reported that mutations in the *FBXL3* gene (C358S or I364T) markedly lengthened the circadian period in mice, and demonstrated that FBXL3 regulates the circadian rhythms by binding to CRY1 and CRY2 and promoting their degradation through the ubiquitin–proteasome pathway [17–19]. The FBXL family comprises 21 proteins with low sequence homology and diverse functions, and its members are often referred to as “orphan proteins”. FBXL3 interacts with SKP1 through its F-box domain to form the SCF complex, which ubiquitinates CRY1 and CRY2, thereby regulating their stability [20–22]. Further studies revealed that FBXL3 and CRY2 cooperate in the degradation of c-MYC, linking circadian regulation to cancer biology [23]. Hirano et al. proposed a model in which FBXL3 localizes to the nucleus while FBXL21 localizes to the cytoplasm, and the two proteins antagonistically regulate CRY protein stability to ensure proper circadian cycling [20]. In humans, mutations in *FBXL3* have been associated with macrocephaly, developmental delay, and intellectual disability [24, 25]. Despite phenotypic differences across species, the molecular mechanism by which FBXL3 regulates the degradation of circadian proteins appears to be evolutionarily conserved.

There are few studies have explored the involvement of FBXL3 in the molecular clock in domestic animals. In sheep, FBXL3 function appears to be evolutionarily conserved; similar to its role in mice, it interacts with CRY1 to promote its ubiquitination and degradation, and exhibits ubiquitous expression across multiple ovine tissues [26]. Notably, a missense mutation in the *FBXL3* gene (Chr10, g.52736358 C>T, Ramb_v2.0) was identified through whole-genome resequencing of 43 domesticated sheep breeds and 17 wild Asian mouflon populations. This mutation, which shows strong selection toward the A allele in domesticated sheep, is putatively associated with reproductive traits [27].

In this study, we performed selective sweep analysis using more than 290 sheep samples representing diverse reproductive phenotypes across the country, and identified strong positive selection at the *FBXL3* mutation site between year-round and seasonally breeding sheep breeds. Notably, the frequency of the G allele was significantly higher in year-round breeders. Evaluation of estrous phenotypes in seasonally breeding sheep further confirmed that individuals carrying this *FBXL3* point mutation exhibited a markedly higher estrus rate during the non-breeding season than their wild-type counterparts. To further investigate the functional consequences of this mutation, we generated homologous point-mutant mice using the CRISPR/Cas9 gene-editing system. Collectively, these findings suggest that the *FBXL3* mutation has undergone positive selection, likely driven by environmental pressures acting on reproductive traits. Mechanistically, we propose that this mutation may alter the binding affinity between FBXL3 and CRY proteins, thereby modulating the expression of specific components of the seasonal reproduction pathway. These molecular changes may influence HPG-axis output and ultimately enable out-of-season breeding in sheep capable of entering estrus during spring and summer.

## 2 Results

### 2.1 *FBXL3* locus was under the selection during sheep domestication

We conducted genome-wide scans for selective sweeps in Hu sheep and 10 Chinese indigenous sheep breeds. It was found that the *FBXL3* gene region on chromosome 10 was under strong selection and displaying a distinct haplotype between year-round estrous and seasonally estrous sheep breeds (Fig 1A and 1B, S1 Fig). Among them, a nonsynonymous mutation site in the coding region of the *FBXL3* gene (Chr10, g.52736358 C>T, ARS-UI_Ramb_v2.0) showed a high *F*_st_ value ranking. Meanwhile, a recent study, using whole-genome resequencing data from 2,420 sheep across 89 breeds worldwide along with environmental and climatic data, also identified the same mutation site through genome-wide association analysis. They found that the frequency of the mutant allele was higher in regions with shorter daylight duration, suggesting that this site may play a regulatory role in the photoperiodic reproduction of sheep [28].

**Fig. 1.**
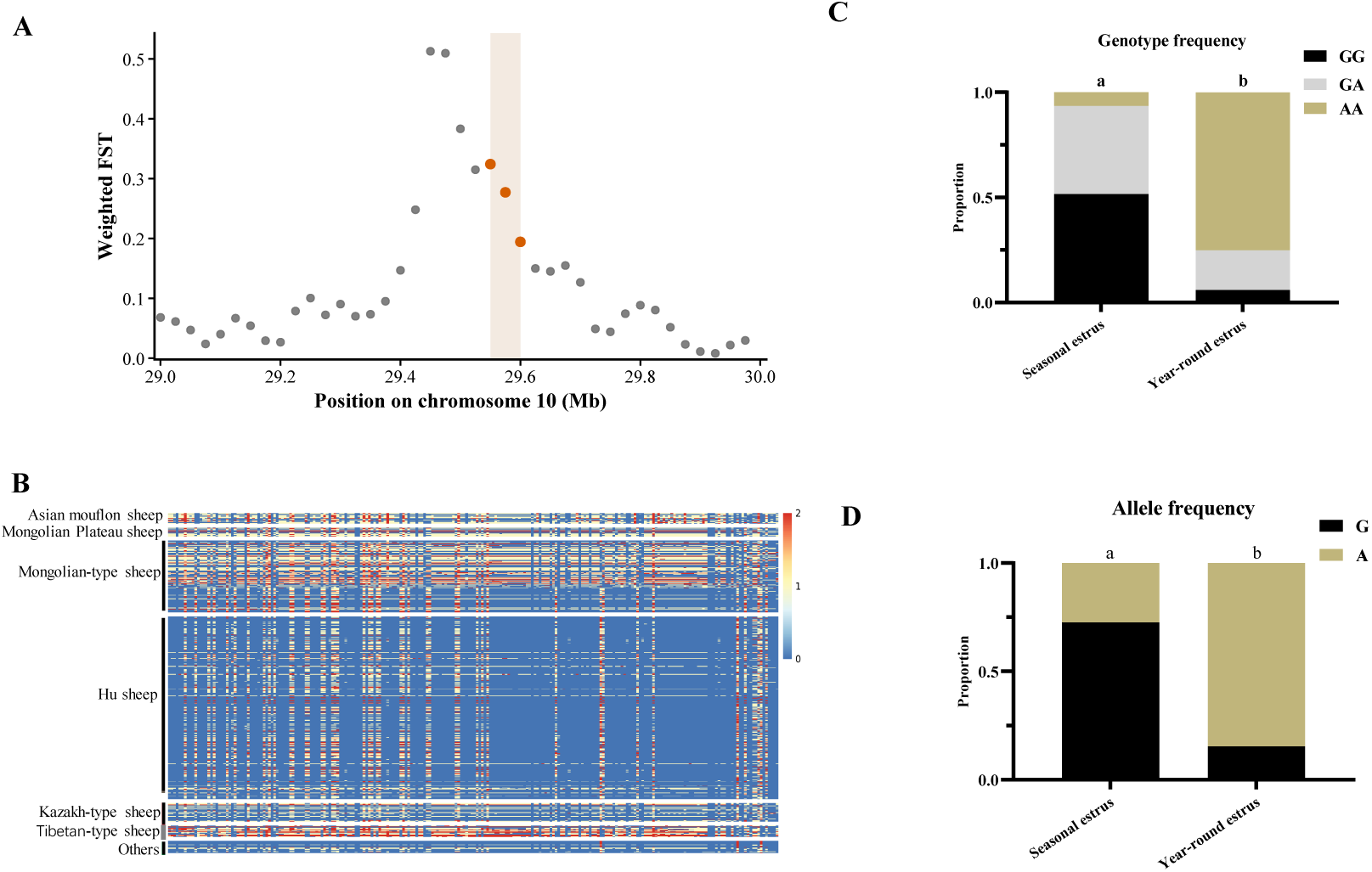
Selection signals at the *FBXL3* locus during sheep domestication. (A) Selective sweep analysis based on whole-genome sequencing data from sheep breeds exhibiting year-round estrus and those showing seasonal breeding patterns. Sliding window size = 50 kb; step size = 25 kb. (B) Haplotypes of the *FBXL3* locus in the Hu sheep population and other sheep breeds. Alleles in Hu sheep and other domestic sheep breeds are indicated in blue and red, respectively. (C) Genotype frequencies of g.52736358G>A in the *FBXL3* gene between year-round-estrous and seasonally breeding sheep breeds. (D) Allele frequencies of g.52736358G>A in the *FBXL3* gene between year-round-estrous and seasonally breeding sheep breeds.

Based on the mapping results above, we selected several breeds with non-seasonal estrus and seasonal estrus traits to genotype the *FBXL3* gene mutation site g.52736358 C>T(ARS-UI_Ramb_v2.0). As shown in Figures 1C and 1D, the genotype and allele frequencies of this site showed highly significant differences between non-seasonal and seasonal estrus breeds (*P* < 0.01). Furthermore, the mutation-derived A allele was the dominant allele in non-seasonal estrus breeds, while the G allele was dominant in seasonal estrus breeds, which is consistent with the findings from previous resequencing analyses.

### 2.2 *FBXL3*^T183M^ was the candidate function mutation

The g.52736358 C>T(ARS-UI_Ramb_v2.0) mutation in the ovine *FBXL3* gene occurs in the exon 3 and results in a missense mutation, substituting threonine (T) with methionine (M) at the 183rd amino acid residue (T183M) of the FBXL3 protein. This mutation site and the corresponding amino acid residue are highly conserved across multiple species (Fig 2A). Using online prediction tools, we assessed potential changes in phosphorylation sites (NetPhos-3.1) and three-dimensional structures (Swiss-Model) associated with the T183M substitution. The results indicated that the phosphorylation site at position 183 was abolished following the mutation, suggesting potential disruption of the functional integrity of this region (Figs 2B and 2C).

**Fig 2.**
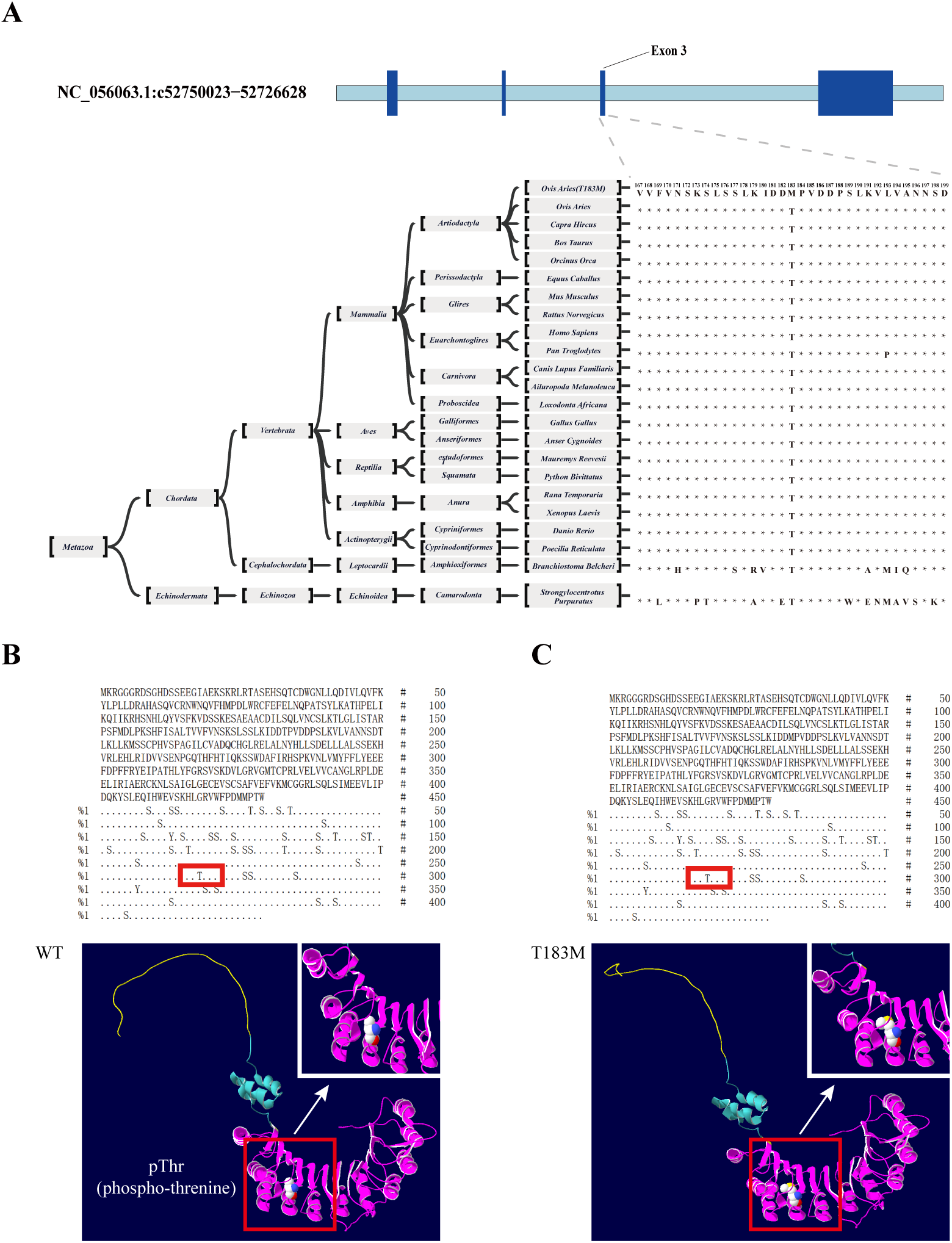
Conservation and protein structure analysis of *FBXL3*^T183M^. (A) Conservative analysis of FBXL3 183rd amino acid of exon3. Predicted three-dimensional structures of FBXL3 in the wild-type (B) and T183M mutant forms (C). FBXL3 protein includes the F-box domain (Aquamarine part in the figure) and the LRR region (Dark Pink part in the figure), where LRR is responsible for recognizing substrate proteins and binding to them, and the T183M mutation is located in the LRR region.

Structurally, threonine possesses a polar, hydrophilic R group, whereas methionine contains a non-polar, hydrophobic R group. The predicted three-dimensional model further suggested that the T183M substitution could increase steric hindrance in the local protein environment, potentially interfering with normal protein–protein interactions (Figs 2B and 2C). Collectively, these alterations may reduce the substrate-binding capacity of the FBXL3 (T183M) variant, impair the function of the SCF^FBXL3^ complex, and consequently slow the rate of ubiquitination and degradation of substrate protein to keep proper circadian cycling.

### 2.3 *FBXL3*^T183M^ sheep showed altered affinity to bind substrate proteins and hormone levels

The above findings suggest that FBXL3 (T183M) may have a reduced capacity to bind substrate proteins. To test whether this mutation affects the interaction between FBXL3 and its substrate CRY1—and whether changes in circadian gene expression could be attributed to altered binding, co-immunoprecipitation (Co-IP) was performed in sheep mutants. *FBXL3* is broadly expressed across tissues, and liver samples are relatively easy to obtain. Therefore, liver tissue was selected for the Co-IP experiment. The results showed that FBXL3 retained the ability to bind CRY1 in sheep, however, the binding affinity of the FBXL3 (T183M) mutant was notably weaker compared to the wild type (Fig 3A).

**Fig. 3.**
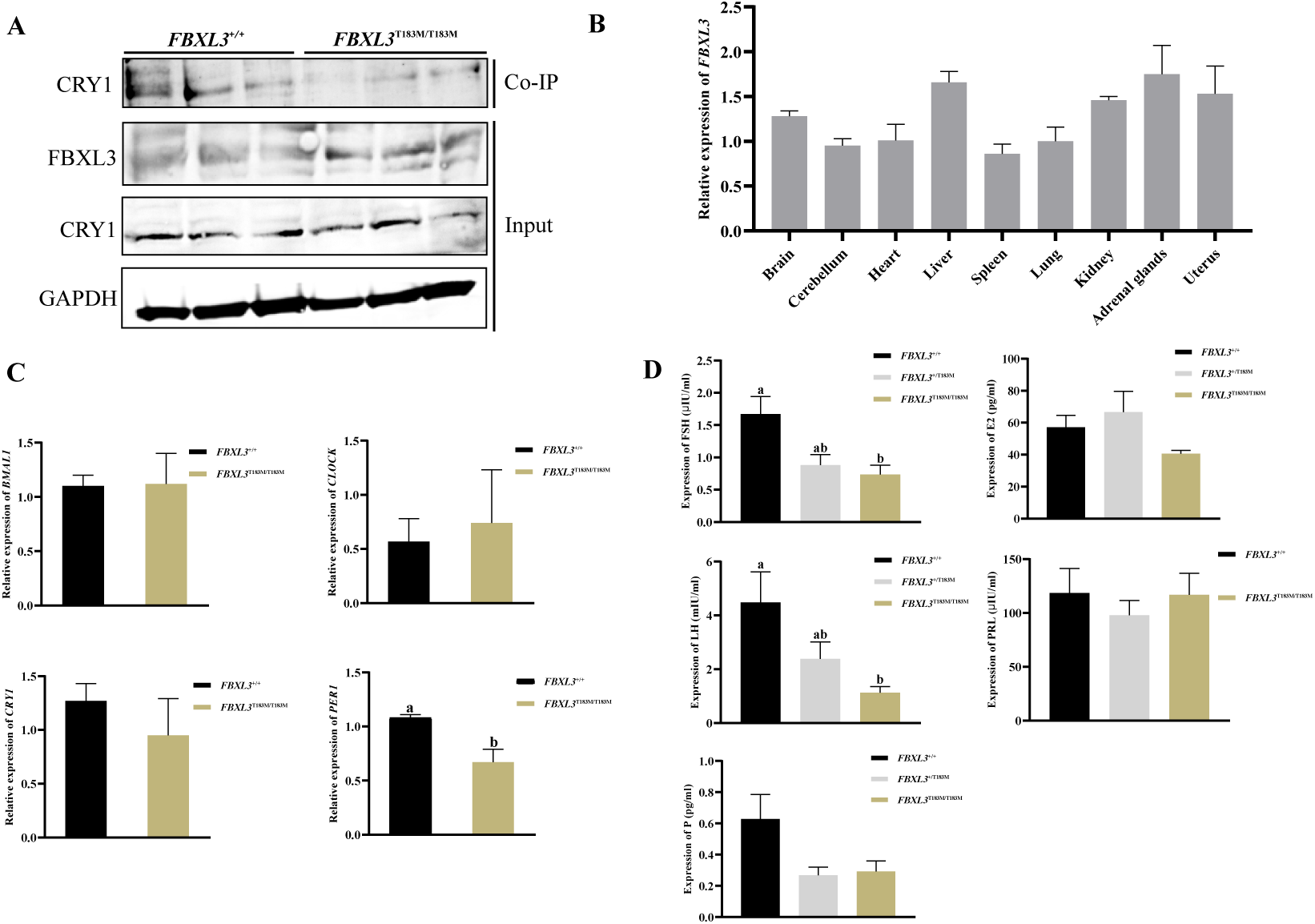
Effects of the FBXL3 T183M mutation on substrate-binding affinity and reproductive hormone levels in sheep. (A) Interaction between FBXL3 and CRY1 proteins in Hu sheep. (B) Tissue expression profile of FBXL3 in Hu sheep. (C) Expression of core circadian clock genes in wild-type and T183M mutant sheep. (D) Reproductive hormone levels in wild-type and T183M mutant sheep, as determined by radioimmunoassay (RIA).

Real-time quantitative PCR further revealed that *FBXL3* is widely expressed across multiple tissues, including the brain, cerebellum, heart, liver, spleen, lung, kidney, adrenal glands, and uterus (Fig 3B). Given that FBXL3 is a critical component of the ubiquitination-mediated degradation of CRY proteins, and that previous studies have shown FBXL3 mutations can delay CRY degradation and thereby lengthen circadian rhythms, the expression of four core circadian rhythm genes were examined. Compared with the wild-type of *FBXL3* with GG genotype, the mutant AA genotype showed decreased expression of *CRY1* and *PER1*, a slight increase in *CLOCK*, and minimal change in *BMAL1* gene expression. Notably, the reduction in *PER1* expression between the GG and AA genotypes was significant (*P* < 0.05) (Fig 3C). These transcriptional changes were also accompanied by genotype-dependent differences in hormone levels, and the hormone levels of ewes with different genotypes during the anestrus period were measured. The secretion levels of FSH and LH were significantly reduced in heterozygous and homozygous mutant individuals compared with the wild type (*P* < 0.05), while the secretion level of progesterone was also lower in heterozygous and homozygous mutants than in the wild type, but the change was not significant. The changes in prolactin and estrogen levels between wild and mutant individuals were not significant (Fig 3D).

### 2.4 *FBXL3*^T183M^ can affect estrus status in sheep

The sheep model with ovariectomized, estradiol-treated ewes and controlled-light conditions was established as previously described [29]. Pituitary tissues from sheep under different photoperiods for real-time quantitative PCR were sampled. The results showed that *FBXL3* expression in the pituitary increased initially and then declined as the photoperiod shifted from short days to long days and with the continued extension of long-day exposure (Fig 4A). This dynamic expression pattern suggests that *FBXL3* may play an important role in regulating seasonal estrus, with temporal changes in its expression potentially affecting gene function. Moreover, as the duration of long days increased, a larger proportion of wild-type sheep entered anestrus, whereas a higher percentage of mutant-type sheep remained in estrus (Fig 4B).

**Fig 4.**
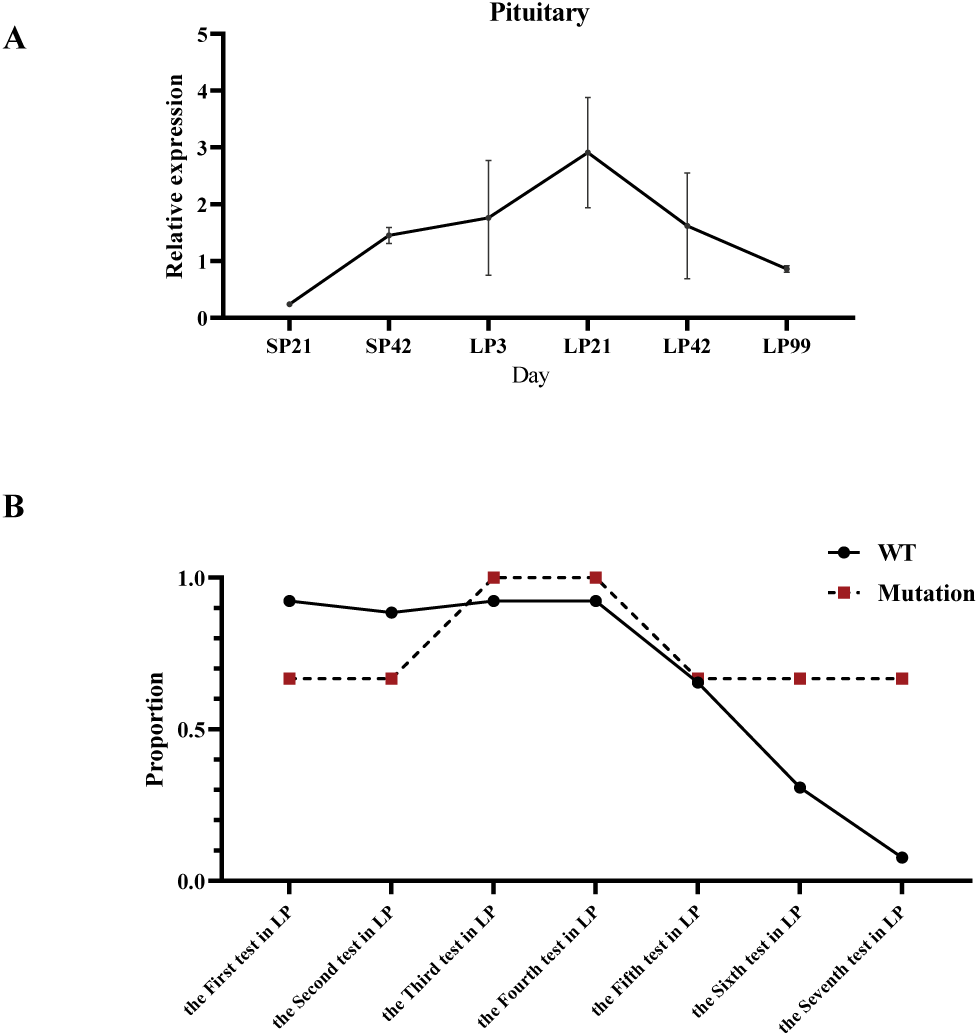
FBXL3 T183M influences estrous status in sheep. (A) Expression of *FBXL3* in the pituitary gland of sheep under different photoperiods (n=3). (B) Estrus rates of sheep under different photoperiods in an ovariectomized and estradiol-implanted (OVX + E_2_) sheep model (WT=26, Mutation=3).

### 2.5 *FBXL3*^T183M^ mouse can prolong circadian rhythms

Given the high conservation of the FBXL3 protein sequence between sheep and mice (97% similarity)(Fig 2A), a mouse model carrying the homologous point mutation (T182M) was generated to evaluate whether the ovine FBXL3 T183M mutation is involved in the regulation of circadian rhythms. The results showed that the T182M mutation significantly prolonged circadian rhythms in mice (Fig 5A, S2 Fig). Compared with wild-type mice, heterozygous mutants exhibited a 0.51 h (*P* < 0.05) longer rhythm, while homozygous mutants showed an extension of 1.00 h (*P* < 0.05)(Fig 5B).

**Fig 5.**
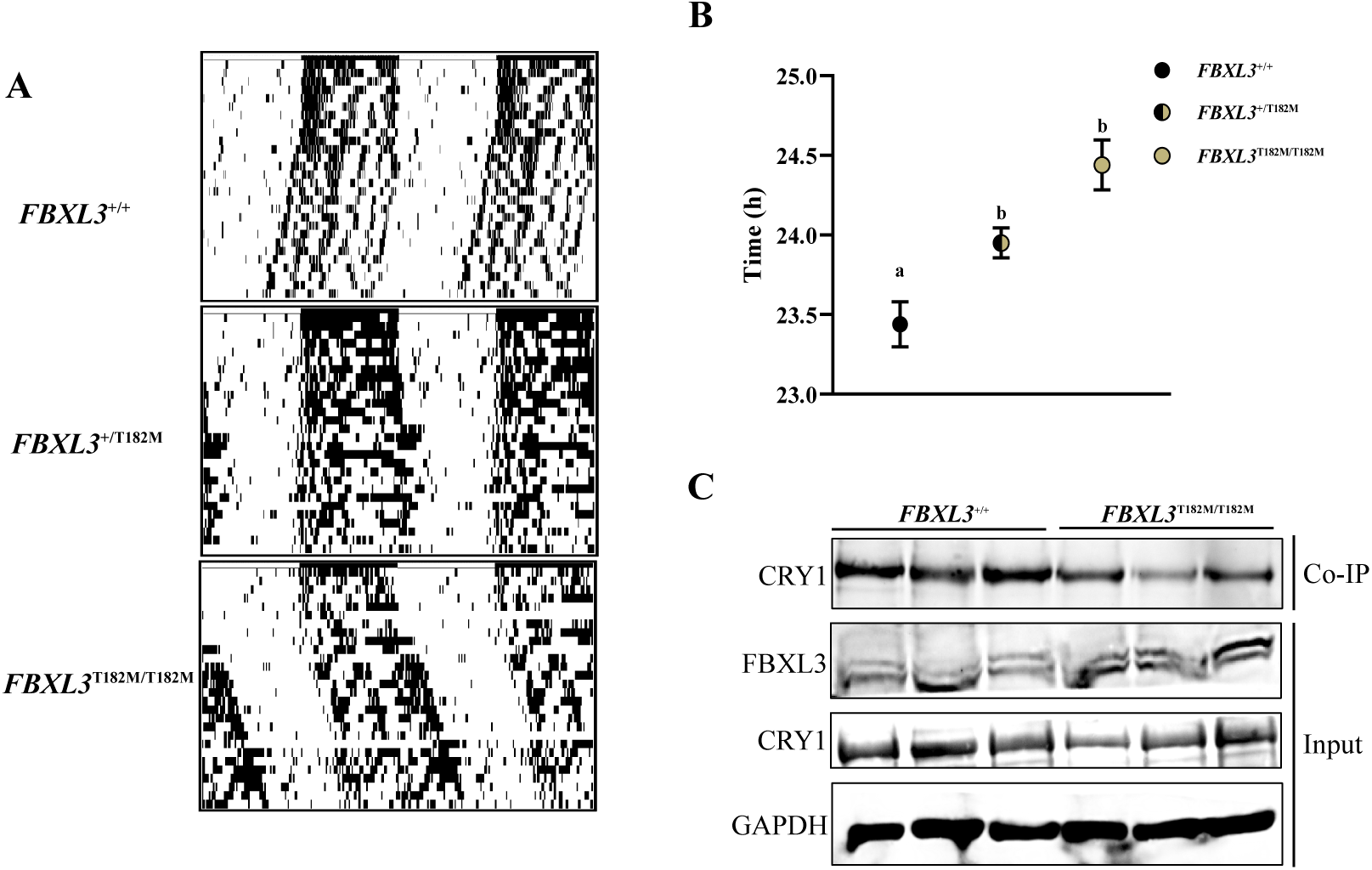
The FBXL3 T183M mutation of FBXL3 protein extends the circadian rhythm in homologous recombination mice. (A) Circadian rhythm map in wild-type and T183M mutant sheep. (B) Circadian rhythm duration of mice with different genotypes. (C) Interaction between FBXL3 and CRY1 proteins in mouse livers.

To further investigate the molecular mechanism, we conducted a Co-IP assay in mice. The results showed that the affinity of FBXL3 (T182M) for CRY1 was markedly reduced compared with the wild type (Fig 5C), suggesting that the T182M mutation impairs the function of FBXL3 within the SKP1-CUL1-F-box (SCF) E3 ubiquitin ligase complex, thereby weakening its ability to promote CRY1 ubiquitination and degradation.

We next examined the effects of this mutation on the expression of core circadian rhythm genes. Quantitative real-time PCR (qRT-PCR) showed that compared with wild-type mice, mutant individuals significantly decreased *CRY1* expression (*P* < 0.05), along with reduced *PER1* expression. Conversely, *CLOCK* and *BMAL1* were upregulated, with *CLOCK* showing a significant increase (*P* < 0.05)(Fig 6A). The results of Western blot showed that the expression of CLOCK protein in mutant individuals increased, while the expressions of CRY1, PER1, and BMAL1 showed no significant changes (Figure 6B, and S3 Fig).

**Fig 6.**
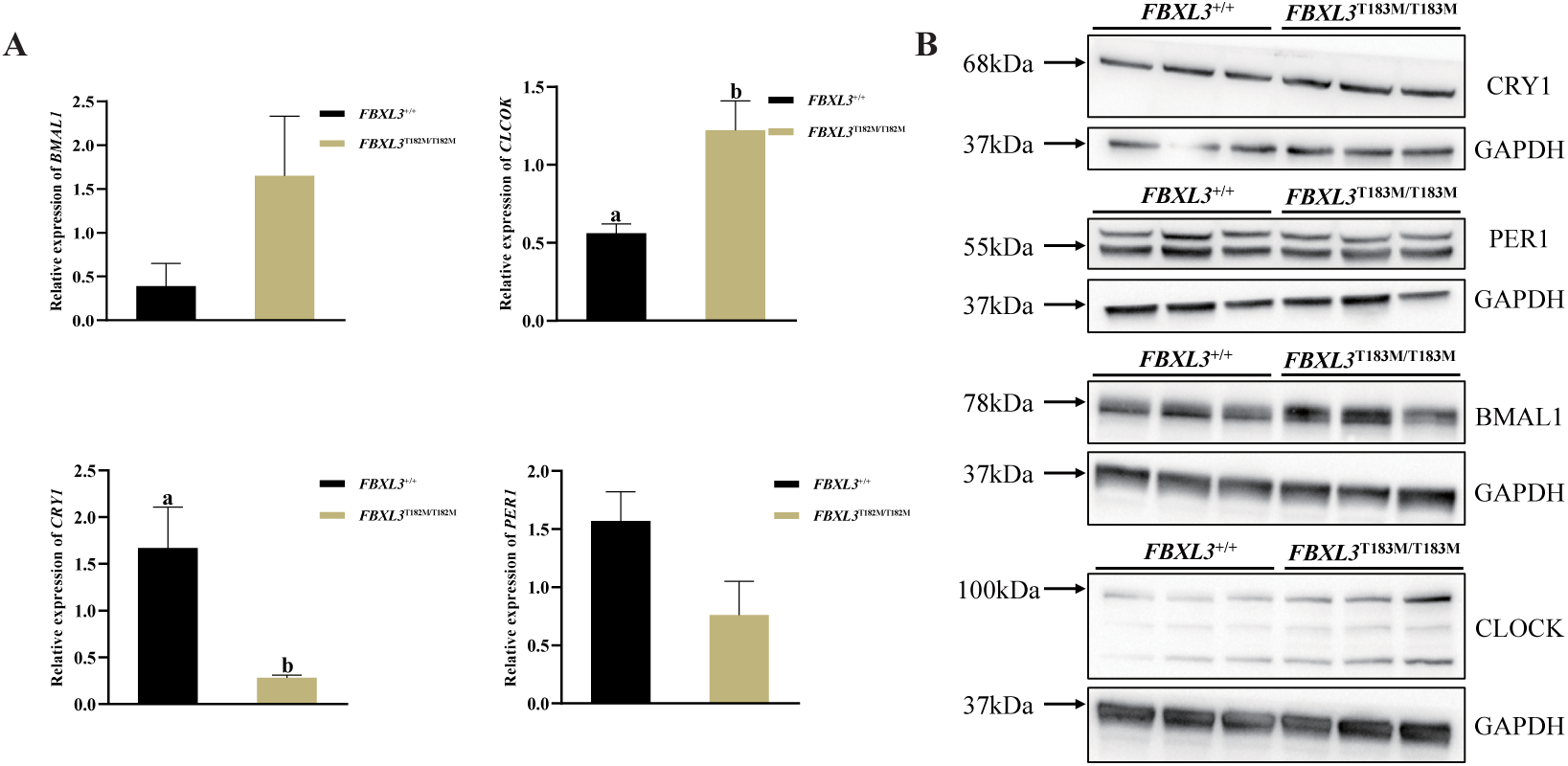
Expression of core circadian clock genes in wild-type and T183M mutant mice. (A) mRNA expression levels of core circadian clock genes, including CRY1, PER1, CLOCK, and BMAL1(qRT-PCR of liver tissue). (B) Protein expression levels of core circadian clock genes, including CRY1, PER1, CLOCK, and BMAL1.

## 3 Discussion

It has long been recognized that domestication and artificial selection can both attenuate reproductive seasonality in livestock. Researchers applied traditional phenotypic selection to a crossbred sheep population for 10 consecutive years, targeting estrus and fertilization during the non-breeding season. Ultimately, the selected lines achieved a fertilization rate of 17% during the seasonal anestrus, whereas the unselected control group exhibited negligible reproductive activity, nearing a zero fertilization rate during the spring [30]. These findings indicate that although estrus-related traits have very low heritability (<0.1), they are still amenable to genetic selection and represent polygenic traits with small individual effects. *FBXL3* is a key component of the circadian clock machinery, and mutations in this gene can disrupt normal molecular clock cycling. *FBXL3* is widely expressed across multiple tissues, underscoring its systemic biological importance. Hugues et al. demonstrated that *ovine FBXL3* is functionally conserved and shows a high degree of homology with its murine counterpart. FBXL3 binds to CRY1 and promotes its ubiquitination and degradation [26]. Notably, Kijas et al. also identified the same missense mutation in the sheep *FBXL3* gene (Chr10, g.52736358 C>T, Ramb_v2.0), suggesting a potential association with estrous traits in sheep [27]. However, the mechanism by which *FBXL3* influences estrous traits through photoperiodic regulation remains unclear.

Numerous studies have shown that FBXL3 and CRY2 cooperatively promote c-MYC degradation, thereby linking circadian clock regulation to cancer biology [23]. Hirano et al. proposed a model in which FBXL3 is predominantly localized in the nucleus, whereas FBXL21 resides mainly in the cytoplasm; through their antagonistic actions, these two proteins jointly regulate CRY protein stability and maintain proper circadian oscillation [20]. In addition, mutations in human FBXL3 have been clinically associated with macrocephaly, developmental delay, and intellectual disability [24, 25]. Although the phenotypic consequences of *FBXL3* variants differ across species, the fundamental mechanism by which FBXL3 regulates the degradation of molecular clock proteins appears to be evolutionarily conserved.

### *FBXL3*^T183M^ may alter secondary structure of mRNA

Previous studies have shown that *FBXL3* is widely expressed across various murine tissues, with peak levels detected in the brain [19]. Similarly, *FBXL3* exhibits a broad expression profile in sheep [26], a pattern consistent with our current findings. Bioinformatic analysis predicted that the g.52736358 C>T mutation (Ramb_v2.0), located in the third exon of *FBXL3*, may alter its mRNA secondary structure. Because mRNA secondary structure is an important determinant of transcript stability, such alterations may lead to tissue-specific differences in gene expression, as previously documented for variants in the MYH7 gene [31]. In the present study, altered mRNA secondary structure, together with insufficient compensatory transcription in response to transcript degradation, may explain the expression differences observed in tissues such as the liver and kidney.

Animal metabolism is governed by both circadian and seasonal rhythms, with the liver and kidney serving as the primary metabolic organs [32]. Previous research demonstrated that the FBXL3 regulates hepatic circadian rhythmicity and glucose homeostasis by promoting CRY1 degradation [33]. Therefore, transcriptional variation in *FBXL3*, together with mutation-induced protein dysfunction, may also contribute to functional differences. Previous studies have reported that under changing photoperiods, specifically during the transition from short-day to long-day conditions, *FBXL3* expression initially increases and then gradually decreases with prolonged long-day exposure. This dynamic expression pattern suggests that *FBXL3* may play an important role in mediating the transition between, as well as the maintenance of, anestrus and estrus phases.

### FBXL3 and Its Role in Circadian Regulation and Seasonal Reproduction

Light variation is the main factor influencing circadian rhythms *in vitro*, whereas *in vivo* rhythmicity is primarily maintained by the dynamic balance between transcriptional activation and repression mediated by clock genes [34, 35]. The core clock components in mammals—CLOCK, BMAL1, PER, and CRY—constitute a transcription–translation feedback loop that drives circadian oscillations [36]. Within this loop, CRY proteins act as potent negative repressors of the circadian master clock, playing an essential role in maintaining rhythmic homeostasis [37, 38]. As a key subunit of the SCF^FBXL3^ ubiquitin ligase complex (SKP1 – CUL1 – F-box protein complex containing the F-box protein FBXL3), FBXL3 utilizers its F-box recruit SCF components and its leucine-rich repeat (LRR) domain to recognizes specific substrates [23]. Acting primarily in the nucleus, the SCF^FBXL3^ complex mediates the ubiquitination and proteasomal degradation of CRY1 and CRY2, thereby fine-tuning circadian rhythmicity in mammals [23, 24, 39]. Research on *FBXL3* began in 2007, when Siepka et al. and Godinho et al. independently identified ENU-induced mutations that extended the circadian period in mice [18, 19]. These mutations altered amino acid residues in FBXL3, impairing SCF^FBXL3^-mediated degradation of CRY1/CRY2 and disrupting the equilibrium between transcriptional activation and repression and establishing a new balance that lengthens the circadian cycle. Beyond the core loop, auxiliary regulators such as Rev-erbα form additional feedback circuits that integrate with the main oscillator, together governing rhythmic transcription of approximately 40% of the genome [38]. Notably, Shi et al. demonstrated that FBXL3 links the core and auxiliary feedback loops through interactions with Rev-erbα, underscoring its pivotal role in circadian regulation [21].

In sheep, FBXL3 exhibits conserved function, promoting CRY degradation similarly to its paralog FBXL21, albeit with partial redundancy [26]. Alternative splice variants of FBXL3 have also been identified, but these isoforms display diminished efficiency in mediating CRY1 degradation. By regulating CRY1/CRY2 turnover, FBXL3 modulates the core circadian feedback loop, driving rhythmic oscillations in PER CRY expression; typically, PER levels peak during the day, whereas CRY levels culminate during the night [40]. Alterations in the phase relationship between PER and CRY, or in their protein–protein interactions under different photoperiods, may provide a molecular basis for photoperiodic responsiveness [2, 40, 41]. Thus, *FBXL3* may influence mammalian seasonal reproduction through modulation of circadian gene expression and phase alignment.

While direct evidence linking *FBXL3* to seasonal reproduction remains limited, its strong association with circadian rhythm regulation suggests a mechanistic connection. Notably, *FBXL3* expression increases and subsequently decreases during prolonged long-day photoperiods, implying potential involvement in the physiological transition between anestrus and estrus. Bioinformatic analysis further indicated that a missense mutation (g.52736358 C>T, Ramb_v2.0) in the *FBXL3* gene leads to loss of phosphorylation at residue 183. Given that phosphorylation is a key post-translational modification regulating protein activity [42], this alteration may impair FBXL3 function. Moreover, genotyping revealed significant differences in both genotype and allele frequencies at the *FBXL3* g.52736358 C>T(Ramb_v2.0) locus between sheep breeds with distinct estrous patterns, suggesting a potential association between *FBXL3* polymorphism and seasonal reproductive behavior.

## 4 Conclusions

In summary, this study identifies a pivotal missense mutation (T183M) in the *FBXL3* gene that is strongly associated with seasonal breeding traits in sheep. By integrating population genetic analysis, physiological modeling, and functional validation in a homologous mutant mouse model, we demonstrate that this mutation perturbs circadian rhythm by weakening the interaction between FBXL3 and CRY1. This molecular disruption lengthens the circadian period and dysregulates the expression of core clock genes. Our findings establish a mechanistic link between the circadian clock system and seasonal reproduction, suggesting that *FBXL3* plays an evolutionarily conserved role in modulating endogenous circannual rhythms. Importantly, the T183M mutation may impair photoperiodic signal transduction, thereby influencing estrous seasonality in sheep. Collectively, this work provides novel insights into the genetic basis of seasonal breeding and identifies a potential molecular target for improving reproductive efficiency in the sheep industry.

## 5 Material and methods

### 5.1 Ethics statement

All animal experiments were performed in accordance with the guidelines of the Animal Advisory Committee at Institute of Genetics and Developmental Biology, Chinese Academy of Sciences (Approval Number: AP2022015-C2).

### 5.2 Experimental animals and genetic data collection

Genetic mapping used Hu sheep as the representative breed for non-seasonal estrus and Mongolian breeds as the representative for seasonal estrus. Sample came from 371 individuals including 10 Chinese native sheep populations. Specifically, 272 Hu sheep were collected from 8 farms (HU, n=272), and Small-tailed Han sheep (STH, n=9). Other Chinese native sheep populations included in this study were: Sishui Fur Sheep (SSS, n=10), Tan Sheep (TAN, n=10), Hulunbuir Sheep (HULUN, n=21), Sunite Sheep (SUNIT, n=10), Prairie-Type Tibetan Sheep (PT, n=9), Oula Tibetan Sheep (OL, n=10), Bayinbuluke Sheep (BY, n=9), Bashbay Sheep (BSB, n=10), and Altay Sheep (ALS, n=10). All the data are available as indicated in publication [43]. Sample data were from public databases. TAN, STH, PT, OL and BY were from SRP066883. HU, SUNIT and HULUN were from PRJCA029248. SSS, BSB, and ALS were from PRJNA624020.

The large population genotyping validation was divided into two groups: the non-seasonal estrus group included Hu sheep (n=696), Small-tailed Han sheep (n=377) Suffolk sheep (n=39), and Dorper sheep (n=30); the seasonal estrus group included Tan sheep (n=166), Hulunbuir sheep (n=180), Prairie-type Tibetan sheep (n=256), and Sunite sheep (n=100). The sheep breeds and genotyping results are shown in Table S1.

### 5.3 Selection signature

The fixation index (*F*_st_) measures the degree of genetic differentiation between populations based on allele frequency differences. Pairwise *F*_st_ values between populations were calculated using VCFtools with a sliding window size of 50 kb and a step size of 25 kb. Genomic regions within the top 0.1% of *F*_st_ values were considered significant candidate loci. To identify loci associated with reproductive seasonality, *F*_st_ was calculated between the year-round estrous group, comprising Hu sheep and STH populations, and the seasonal estrous group, comprising the TAN, SUNIT, PT, OL, HULUN, BY, SSS, ALS, and BSB populations. The top 1‰ significant loci and annotation results are shown in S2 Table.

### 5.4 Sheep estrous cycle determination

*FBXL3* gene expression detection was performed on six stages, with three wild-type animals per stage. Estrus data measurements included 26 wild-type and 3 mutant-type Sunite sheep. All animals were housed at the breeding base of the Tianjin Animal Husbandry and Veterinary Research Institute. First, we herded all the sheep into the light-controlled room and set the lighting to a short photoperiod (SP), with that day recorded as SP0. On SP25, we prepared the ovariectomized and estradiol-implanted (OVX + E_2_) ewe model and the light-controlled room as previously described [44]. The short-day photoperiod (light : darkness = 8h:16h) and long-day photoperiod (light:darkness = 16h:8h) conditions were used as indicated in previous study [29]. Afterwards, the ewes’ estrus status was determined using the ram test. The day after all the ewes showed estrus, the short photoperiod was changed to a long photoperiod, with that day recorded as LP 0. The estrous cycle of Sunite sheep is approximately 15–19 days; based on this, we conducted a daily ram test for all ewes starting before each ewe’s natural estrus until estrus was detected in all ewes, defining this as one natural estrous cycle. This cycle was repeated until the ewes entered anestrus. During this process, we recorded whether each ewe exhibited estrus. Day 1 of the short-day photoperiod was SP1, and day 1 of the long-day photoperiod was LP1, with sampling points on SP21, SP42, LP3, LP21, LP42, and LP99.

### 5.5 Protein structure prediction

FBXL3 protein sequences from various species mentioned in the text were downloaded from NCBI, including the sheep sequence (accession number: NP_001123211). The multi-species FBXL3 protein sequence alignment was performed using MEGA-X software. Protein phosphorylation site prediction was conducted using the online tool NetPhos 3.1 (https://services.healthtech.dtu.dk/services/NetPhos-3.1/); protein structure prediction before and after mutation was performed using the online tool SWISS-MODEL (https://swissmodel.expasy.org/), and the differences were compared using Swiss-PDBViewer.

### 5.6 Quantitative Real-Time PCR

Tissue samples were collected from three wild-type and three mutant-type healthy 2-3-year-old Sunite ewes at the same time points for different genotypes, with tissue samples quickly frozen in liquid nitrogen and transported on dry ice. According to the reference sequences Primers of Sheep FBXL3 (GenBank accession number: NM_001129739.1), CRY1 (accession number: NM_001129735.1), PER1 (accession number: XM_027974927.2), CLOCK (accession number: NM_001130932.1), BMAL1 (accession number: NM_001129734.1) and mouse CRY1 (GenBank accession number: NM_007771.3), PER1 (accession number: NM_001159367.2), CLOCK (accession number: NM_001289826.1), and BMAL1 (accession number: NM_001243048.1) gene sequences were designed based on the sheep reference genome (ARS-UI Ramb v3.0) and GRCm39(GCF_000001635.27) at the NCBI (National Center of Biotechnology Information). qPCR was performed using Taq Pro Universal SYBR qPCR Master Mix (Q712-03, Vazyme) in a 10 μL reaction volume containing 0.3 μL each of forward and reverse primers, 1 μL of cDNA (50 ng), 5 μL of qPCR mix, and 3.4 μL of ddH_2_O. Amplificaiton conditions were:95 °C for 1 min; 40 cycles of 95 °C for 10 s, 60 °C for 15 s, and 72 °C for 15 s.β-actin was used as the endogenous control, and relative expression levels were calculated using the 2^−^^(ΔΔCt) method. Primer sequences are detailed in S3 Table.

### 5.7 Generation of a point mutation knock-in mouse model

For FBXL3 protein, the 183rd amino acid (T183, codon ACG) in sheep (ENSOARG00000016056) which corresponds to the 182th amino acid (M182, codon ACC) in mouse(ENSMUSG00000022124) was conserved. The Fbxl3 point mutation (p.T182M) mouse model was established on a C57BL/6N background via CRISPR/Cas9-mediated genome editing (Biocytogen Pharmaceuticals, Beijing). To introduce the specific mutation, a donor vector containing the p.T182M substitution and flanking homology arms was designed to repair the DSBs induced by the Cas9/sgRNA complex. One-cell stage embryos (C57BL/6N) were co-injected with Cas9 mRNA, sgRNA, and the donor template, followed by embryo transfer into pseudopregnant ICR foster mothers.

Genomic DNA was extracted from tail biopsies of F0 mice. The target region was amplified by PCR and subjected to Sanger sequencing to confirm the presence of the intended point mutation. Positive founders were further bred with wild-type C57BL/6J mice to establish germline transmission.

### 5.8 Rhythm measurement

The rhythm measurement instrument was the LE 3806 MULTICOUNTER from Panlab (Barcelona, Spain), and the data collection software was SEDACOM_v2001. Bedding, food, and water were added to the mouse cages, and the mice were placed inside. The device was activated to record for 7 days under light:dark conditions of 12h:12h to acclimate the mice to the testing environment. On the 8th day, external lights were turned off, creating a 24-hour dark environment to minimize external light exposure. Rhythm measurement continued for more than three weeks in complete darkness. The results are shown in S4 Table, expressed as mean ± SEM.

### 5.9 Co-immunoprecipitation and immunoblot analysis

For co-immunoprecipitation (Co-IP) experiments, the sheep livers were lysed in 1 mL of RIPA buffer (50 mM Tris-HCl, pH 8.0; 150 mM NaCl; 0.5 mM EDTA, 0.1% SDS, 1% sodium deoxycholate, 1% Triton X-100 supplemented with protease inhibitor PMSF) , after homogenization with a tissue homogenizer, centrifuged at 12,000 g at 4°C for 10 min. measure the protein concentration of the supernatant using the BCA method after extraction. Subsequently, the Co-IP experiment was performed according to the PK0007 (Proteintech) protocol, after adding the FBXL3 primary antibody, the sample was incubated overnight at 4°C. For immunoblot analyses, the samples were fractionated by sodium dodecyl sulfate polyacrylamide gel electrophoresis (SDS-PAGE) and transferred onto polyvinylidene fluoride membranes (Merck). The membranes were first blocked with 5% (w/v) non-fat milk in TBST, and subsequent immunoblot analysis was performed with the indicated primary antibodies, followed by horseradish peroxidase (HRP)-conjugated secondary antibodies The protein bands were visualized using Pierce^TM^ ECL Western Blotting Substrate (Thermo Scientific) according to the manufacturer’s instructions. Results were analyzed using ImageJ for gray value analysis. The antibodies are shown in S5 Table.

### 5.10 Radioimmunoassay (RIA)

Radioimmunoassay was conducted by the Beijing North Biotechnology Research Institute. The samples are Hu sheep serum, comprising 3 wild-type, 4 heterozygous, and 3 mutant individuals, transported entirely on dry ice. Measurement of hormones related to reproduction, including prolactin (PRL), follicle-stimulating hormone (FSH), luteinizing hormone (LH), estradiol (E2), and progesterone. The antibodies are shown in S5 Table. A specific antibody is incubated with a known amount of radiolabeled antigen and the sample or standard containing unlabeled antigen. The labeled and unlabeled antigens compete for limited antibody binding sites until equilibrium is reached. Subsequently, the antibody-bound fraction is separated from the free antigen, and the radioactivity of one fraction is measured using a gamma counter. The concentration of the target analyte in the sample is then determined by comparison with a standard curve generated from known concentrations. The results are shown in S6 Table.

## Supporting information

S1 Fig. Selective sweep signals between year-round estrous and seasonally estrous sheep breeds.

(PDF)

S2 Fig. Experimental graphs of the circadian rhythm activity of mice.

(PDF)

S3Fig. Data quantification of panel Fig 6B.

(PDF)

S1 Table. The number of three genotypes in different sheep breeds.

(XLSX)

S2 Table. Genome-wide top 1‰ significant loci and their annotations identified by selective sweep analysis between year-round estrous and seasonally estrous sheep breeds.

(XLSX)

S3 Table. Primers for Quantitative PCR.

(XLSX)

S4 Table. Rhythmic Duration in Mice with Different FBXL3 Genotypes.

(XLSX)

S5 Table. The number of three genotypes in different sheep breeds.

(XLSX)

S6 Table. Differential expression of reproductive hormones in sheep with different FBXL3 genotypes.

(XLSX)

## Acknowledgments

We thank the support of the Xihe high-performance computing platform of the National Research Facility for Phenotypic and Genotypic Analysis of Model Animals (Beijing), China Agricultural University.

## Author contributions

Conceptualization: Ran Li, Qiuyue Liu.

Data curation: Tingting Li, Haitao Wang.

Formal analysis: Yang Yang, Na Zhang, Xinxiao Jing.

Funding acquisition: Mingxing Chu, Qiuyue Liu.

Investigation: Yang Yang, Na Zhang, Tingting Li, Haitao Wang, Haoran Zhang, Ran Di, Qing Xia.

Project administration: Mingxing Chu, Qiuyue Liu.

Resources: Xun Huang, Runlin Ma.

Methodology: Yang Yang, Na Zhang, Xiaoyun He, Xiaofei Guo, Xiaosheng Zhang, Yu Jiang.

Supervision: Qiuyue Liu.

Writing – original draft: Yang Yang, Na Zhang.

Writing – review and editing: Yang Yang, Na Zhang, Ran Li, Mingxing Chu, Qiuyue Liu.

## References

1. Ikegami K, Yoshimura T. The hypothalamic-pituitary-thyroid axis and biological rhythms: The discovery of TSH’s unexpected role using animal models. Best Practice & Research Clinical Endocrinology & Metabolism. 2017;31(5):475–85.

2. Ikegami K, Yoshimura T. Circadian clocks and the measurement of daylength in seasonal reproduction. Mol Cell Endocrinol. 2012;349(1):76–81. doi: 10.1016/j.mce.2011.06.040. PMID: 21767603.

3. Ralph MR, Foster RG, Davis FC, Menaker M. Transplanted suprachiasmatic nucleus determines circadian period. Science. 1990;247(4945):975–8. doi: 10.1126/science.2305266. PMID: 2305266.

4. Xu P, Berto S, Kulkarni A, Jeong B, Joseph C, Cox KH, et al. NPAS4 regulates the transcriptional response of the suprachiasmatic nucleus to light and circadian behavior. Neuron. 2021;109(20):3268–82.e6. doi: 10.1016/j.neuron.2021.07.026. PMID: 34416169.

5. Brainard GC, Hanifin JP, Greeson JM, Byrne B, Glickman G, Gerner E, et al. Action spectrum for melatonin regulation in humans: evidence for a novel circadian photoreceptor. The Journal of neuroscience : the official journal of the Society for Neuroscience. 2001;21(16):6405–12. doi: 10.1523/jneurosci.21-16-06405.2001. PMID: 11487664.

6. Pandi-Perumal SR, Srinivasan V, Maestroni GJ, Cardinali DP, Poeggeler B, Hardeland R. Melatonin: Nature’s most versatile biological signal? The FEBS journal. 2006;273(13):2813–38. doi: 10.1111/j.1742-4658.2006.05322.x. PMID: 16817850.

7. Johnston JD, Tournier BB, Andersson H, Masson-Pévet M, Lincoln GA, Hazlerigg DG. Multiple effects of melatonin on rhythmic clock gene expression in the mammalian pars tuberalis. Endocrinology. 2006;147(2):959–65. doi: 10.1210/en.2005-1100. PMID: 16269454.

8. von Gall C, Garabette ML, Kell CA, Frenzel S, Dehghani F, Schumm-Draeger PM, et al. Rhythmic gene expression in pituitary depends on heterologous sensitization by the neurohormone melatonin. Nat Neurosci. 2002;5(3):234–8. doi: 10.1038/nn806. PMID: 11836530.

9. Dardente H, Wyse CA, Birnie MJ, Dupré SM, Loudon AS, Lincoln GA, et al. A molecular switch for photoperiod responsiveness in mammals. Current Biology Cb. 2010;20(24):2193.

10. Masumoto KH, Ukai-Tadenuma M, Kasukawa T, Nagano M, Uno KD, Tsujino K, et al. Acute induction of Eya3 by late-night light stimulation triggers TSHβ expression in photoperiodism. Curr Biol. 2010;20(24):2199–206. doi: 10.1016/j.cub.2010.11.038. PMID: 21129973.

11. Dardente H, Lomet D, Robert V, Decourt C, Beltramo M, Pellicer-Rubio MT. Seasonal breeding in mammals: From basic science to applications and back. Theriogenology. 2016;86(1):324–32. doi: 10.1016/j.theriogenology.2016.04.045. PMID: 27173960.

12. Hanon EA, Lincoln GA, Fustin JM, Dardente H, Masson-Pévet M, Morgan PJ, et al. Ancestral TSH mechanism signals summer in a photoperiodic mammal. Curr Biol. 2008;18(15):1147–52. doi: 10.1016/j.cub.2008.06.076. PMID: 18674911.

13. Ono H, Hoshino Y, Yasuo S, Watanabe M, Nakane Y, Murai A, et al. Involvement of thyrotropin in photoperiodic signal transduction in mice. Proceedings of the National Academy of Sciences of the United States of America. 2008;105(47):18238–42. doi: 10.1073/pnas.0808952105. PMID: 19015516.

14. Dupré SM, Miedzinska K, Duval CV, Yu L, Goodman RL, Lincoln GA, et al. Identification of Eya3 and TAC1 as long-day signals in the sheep pituitary. Curr Biol. 2010;20(9):829–35. doi: 10.1016/j.cub.2010.02.066. PMID: 20434341.

15. Lomet D, Cognié J, Chesneau D, Dubois E, Hazlerigg D, Dardente H. The impact of thyroid hormone in seasonal breeding has a restricted transcriptional signature. Cellular and molecular life sciences : CMLS. 2018;75(5):905–19. doi: 10.1007/s00018-017-2667-x. PMID: 28975373.

16. Wood SH, Hindle MM, Mizoro Y, Cheng Y, Saer BRC, Miedzinska K, et al. Circadian clock mechanism driving mammalian photoperiodism. Nature communications. 2020;11(1):4291. doi: 10.1038/s41467-020-18061-z. PMID: 32855407.

17. Busino L, Bassermann F, Maiolica A, Lee C, Nolan PM, Godinho SI, et al. SCFFbxl3 controls the oscillation of the circadian clock by directing the degradation of cryptochrome proteins. Science. 2007;316(5826):900–4. doi: 10.1126/science.1141194. PMID: 17463251.

18. Godinho SI, Maywood ES, Shaw L, Tucci V, Barnard AR, Busino L, et al. The after-hours mutant reveals a role for Fbxl3 in determining mammalian circadian period. Science. 2007;316(5826):897–900. doi: 10.1126/science.1141138. PMID: 17463252.

19. Siepka SM, Yoo SH, Park J, Song W, Kumar V, Hu Y, et al. Circadian mutant Overtime reveals F-box protein FBXL3 regulation of cryptochrome and period gene expression. Cell. 2007;129(5):1011–23. doi: 10.1016/j.cell.2007.04.030. PMID: 17462724.

20. Hirano A, Yumimoto K, Tsunematsu R, Matsumoto M, Oyama M, Kozuka-Hata H, et al. FBXL21 regulates oscillation of the circadian clock through ubiquitination and stabilization of cryptochromes. Cell. 2013;152(5):1106–18. doi: 10.1016/j.cell.2013.01.054. PMID: 23452856.

21. Shi G, Xing L, Liu Z, Qu Z, Wu X, Dong Z, et al. Dual roles of FBXL3 in the mammalian circadian feedback loops are important for period determination and robustness of the clock. Proceedings of the National Academy of Sciences of the United States of America. 2013;110(12):4750–5. doi: 10.1073/pnas.1302560110. PMID: 23471982.

22. Yoo SH, Mohawk JA, Siepka SM, Shan Y, Huh SK, Hong HK, et al. Competing E3 ubiquitin ligases govern circadian periodicity by degradation of CRY in nucleus and cytoplasm. Cell. 2013;152(5):1091–105. doi: 10.1016/j.cell.2013.01.055. PMID: 23452855.

23. Huber AL, Papp SJ, Chan AB, Henriksson E, Jordan SD, Kriebs A, et al. CRY2 and FBXL3 Cooperatively Degrade c-MYC. Molecular cell. 2016;64(4):774–89. doi: 10.1016/j.molcel.2016.10.012. PMID: 27840026.

24. Ansar M, Paracha SA, Serretti A, Sarwar MT, Khan J, Ranza E, et al. Biallelic variants in FBXL3 cause intellectual disability, delayed motor development and short stature. Hum Mol Genet. 2019;28(6):972–9. doi: 10.1093/hmg/ddy406. PMID: 30481285.

25. Confino S, Dor T, Tovin A, Wexler Y, Ben-Moshe Livne Z, Kolker M, et al. A Zebrafish Model for a Rare Genetic Disease Reveals a Conserved Role for FBXL3 in the Circadian Clock System. Int J Mol Sci. 2022;23(4). doi: 10.3390/ijms23042373. PMID: 35216494.

26. Dardente H, Mendoza J, Fustin JM, Challet E, Hazlerigg DG. Implication of the F-Box Protein FBXL21 in circadian pacemaker function in mammals. PLoS One. 2008;3(10):e3530. doi: 10.1371/journal.pone.0003530. PMID: 18953409.

27. Naval-Sanchez M, Nguyen Q, McWilliam S, Porto-Neto LR, Tellam R, Vuocolo T, et al. Sheep genome functional annotation reveals proximal regulatory elements contributed to the evolution of modern breeds. Nature communications. 2018;9(1):859. doi: 10.1038/s41467-017-02809-1. PMID: 29491421.

28. Kang Y. Genetic Characteristics of Sheep Adaptation to Sunshine and Temperature Based on Wholegenome Resequencing. M.Agr. Thesis, Northwest A&F University. 2024. Available from: https://link.cnki.net/doi/10.27409/d.cnki.gxbnu.2024.000565

29. He X, Di R, Wang X, Guo X, Zhang X, Zhang J, et al. Photoperiodic responsiveness in the DNA methylation and gene expression in the hypothalamus of ovariectomized and estradiol-treated ewes. BMC genomics. 2025;26(1):873. doi: 10.1186/s12864-025-12066-y. PMID: 41023860.

30. Notter DR. Genetic Aspects of Reproduction in Sheep. Reproduction in Domestic Animals. 2008;43(s2):122–8.

31. Montag J, Petersen B, Fl?Gel AK, Becker E, Lucas-Hahn A, Cost GJ, et al. Successful knock-in of Hypertrophic Cardiomyopathy-mutation R723G into the MYH7 gene mimics HCM pathology in pigs. 2018;8(1):144–52.

32. Helfer G, Barrett P, Morgan PJ. A unifying hypothesis for control of body weight and reproduction in seasonally breeding mammals. J Neuroendocrinol. 2019;31(3):e12680. doi: 10.1111/jne.12680. PMID: 30585661.

33. Liu Z, Qian M, Tang X, Hu W, Sun S, Li G, et al. SIRT7 couples light-driven body temperature cues to hepatic circadian phase coherence and gluconeogenesis. Nature metabolism. 2019;1(11):1141–56. doi: 10.1038/s42255-019-0136-6. PMID: 32694864.

34. Julius AA, Yin J, Wen JT. Time optimal entrainment control for circadian rhythm. PLoS One. 2019;14(12):e0225988. doi: 10.1371/journal.pone.0225988. PMID: 31851723.

35. Sato TK, Yamada RG, Ukai H, Baggs JE, Miraglia LJ, Kobayashi TJ, et al. Feedback repression is required for mammalian circadian clock function. Nature genetics. 2006;38(3):312–9. doi: 10.1038/ng1745. PMID: 16474406.

36. Mohawk JA, Green CB, Takahashi JS. Central and peripheral circadian clocks in mammals. Annu Rev Neurosci. 2012;35:445–62. doi: 10.1146/annurev-neuro-060909-153128. PMID: 22483041.

37. Anand SN, Maywood ES, Chesham JE, Joynson G, Banks GT, Hastings MH, et al. Distinct and separable roles for endogenous CRY1 and CRY2 within the circadian molecular clockwork of the suprachiasmatic nucleus, as revealed by the Fbxl3(Afh) mutation. The Journal of neuroscience : the official journal of the Society for Neuroscience. 2013;33(17):7145–53. doi: 10.1523/jneurosci.4950-12.2013. PMID: 23616524.

38. Parico GCG, Partch CL. The tail of cryptochromes: an intrinsically disordered cog within the mammalian circadian clock. Cell communication and signaling : CCS. 2020;18(1):182. doi: 10.1186/s12964-020-00665-z. PMID: 33198762.

39. Saran AR, Kalinowska D, Oh S, Janknecht R, DiTacchio L. JMJD5 links CRY1 function and proteasomal degradation. PLoS biology. 2018;16(11):e2006145. doi: 10.1371/journal.pbio.2006145. PMID: 30500822.

40. Lincoln G, Messager S, Andersson H, Hazlerigg D. Temporal expression of seven clock genes in the suprachiasmatic nucleus and the pars tuberalis of the sheep: evidence for an internal coincidence timer. Proceedings of the National Academy of Sciences of the United States of America. 2002;99(21):13890–5. doi: 10.1073/pnas.212517599. PMID: 12374857.

41. Pittendrigh CS. Circadian surfaces and the diversity of possible roles of circadian organization in photoperiodic induction. Proceedings of the National Academy of Sciences of the United States of America. 1972;69(9):2734–7. doi: 10.1073/pnas.69.9.2734. PMID: 4506793.

42. Kosako H, Nagano K. Quantitative phosphoproteomics strategies for understanding protein kinase-mediated signal transduction pathways. Expert review of proteomics. 2011;8(1):81–94. doi: 10.1586/epr.10.104. PMID: 21329429.

43. Liu Z, Zhang N, Wen Z, Wang H, Li T, Ma R, et al. Genomic Insights into the Origin, High Fecundity and Environmental Adaptation of Hu Sheep. Adv Sci (Weinh). 2025;12(37):e06492. doi: 10.1002/advs.202506492. PMID: 40658082.

44. Di R, He J, Song S, Tian D, Liu Q, Liang X, et al. Characterization and comparative profiling of ovarian microRNAs during ovine anestrus and the breeding season. BMC genomics. 2014;15(1):899. doi: 10.1186/1471-2164-15-899. PMID: 25318541.

